# A computational platform for high-throughput analysis of RNA sequences and modifications by mass spectrometry

**DOI:** 10.1101/501668

**Authors:** Samuel Wein, Byron Andrews, Timo Sachsenberg, Helena Santos-Rosa, Oliver Kohlbacher, Tony Kouzarides, Benjamin A. Garcia, Hendrik Weisser

## Abstract

The field of epitranscriptomics is growing in importance, with chemical modification of RNA being associated with a wide variety of biological phenomena. A pivotal challenge in this area is the identification of modified RNA residues within their sequence contexts. Next-generation sequencing approaches are generally unable to capture modifications, although workarounds for some epigenetic marks exist. Mass spectrometry (MS) offers a comprehensive solution by using analogous approaches to shotgun proteomics. However, software support for the analysis of RNA MS data is inadequate at present and does not allow high-throughput processing. In particular, existing software solutions lack the raw performance and statistical grounding to efficiently handle the large variety of modifications present on RNA. We present a free and open-source database search engine for RNA MS data, called NucleicAcidSearchEngine (NASE), that addresses these shortcomings. We demonstrate the capability of NASE to reliably identify a wide range of modified RNA sequences in three original datasets of varying complexity. In a human tRNA sample, we characterize over 20 different modification types simultaneously and find many cases of incomplete modification.

## Introduction

RNA is an extensively modified biological macromolecule. Over 150 chemically distinct modifications have been reported. The presence of methylated adenine, cytosine, and guanine in RNA was uncovered in the 1960s^1^, and pseudouridine has been referred to as the “fifth base” for decades^2^. However, widespread interest in these epitranscriptomic marks has been raised by recent reports that underscore their importance in a wide variety of developmental signalling. In stem cells the intracellular effector proteins SMAD2 and SMAD3 promote binding of the N6-methyladenosine (m6A) writer complex to a subset of mRNAs associated with early cell fate decisions^3^. Likewise, a number of modifications are associated with disease. It has been demonstrated that the loss of taurine modification in the anticodon of mitochondrial tRNA-Leu is responsible for mitochondrial myopathy, encephalopathy, lactic acidosis, and stroke-like episodes (MELAS)^4^. m6A is implicated in obesity^5^ and associated with defects in functional axon regeneration in mice^6^. Aberrant methylation of cytosine-5 (m5C) in tRNAs has been linked to neuro-developmental disorders^7^.

Recent interest in epitranscriptomics has also been spurred by technical advances in next-generation sequencing (NGS) technology, which has allowed modifications in mRNA to be profiled individually. All of the approaches based on Solexa/Illumina sequencing use antibodies to immunoprecipitate modified RNA, and/or apply chemical treatments to alter it and read out modifications as mutations or truncations in the preparation of cDNA^8,9^. The primary caveat of these methods is that only a single type of modification can be profiled in each experiment, and specific chemical and/or antibody reagents do not exist for every modification. Further complications can be caused by lack of specificity of the existing antibodies, in particular m6A and m6Am^10^. Steps have been made towards uncovering modifications directly using long-read sequencing platforms^11,12^, but many technical challenges stand between these approaches and routine use, not least a significant error rate of greater than 13% per single RNA read^13^. NGS-based methods have also generated conflicting results in the past^14,15^, underscoring the need for orthogonal approaches.

Mass spectrometry (MS) is currently the only technique that can directly and comprehensively characterize chemical modifications in RNA sequences. The majority of RNA MS has focused on reducing the RNA to mono-nucleosides and applying workflows analogous to metabolite analysis^16^. While these techniques are effective in determining what modifications are present in a sample, all information about the location and co-occurrence of modifications is lost. This information is critical in complex samples to allow attributing modifications to specific RNAs. Even in simpler cases, modification location and co-occurrence may be important for a phenotypic effect; for example, in microRNA 2’-O-methylation of the 3’-most nucleic acid sterically inhibits 3’ exonuclease digestion, which increases the half-life of the modified microRNA in the cell. For this reason there is interest in analyzing samples in as close to their native states as possible. Analysis of intact RNA oligonucleotides by tandem mass spectrometry (MS/MS) is capable of determining modification sites with single-nucleotide resolution, by comparing mass spectra with a sequence database^17^. However, oligonucleotides are challenging to separate via liquid chromatography (LC) that is compatible with mass spectrometry. The current approach of choice is reversed-phase ion-pair liquid chromatography^18^.

In addition to the experimental challenges, difficulties emerge in interpreting the acquired data. Considerable efforts towards automating data analysis have been made in recent years, starting with SOS^19^ in 2002, Ariadne^20^ in 2009, OMA/OPA^21^ in 2012, and RNAModMapper^22^ in 2017, all of which are programs for database-matching or decoding the complicated patterns of oligonucleotide fragmentation. However, none of the existing software solutions offers key features necessary to analyze data from large-scale experiments. First, no software can efficiently handle the analysis of RNA oligonucleotide data – especially of more complex samples or involving many different modifications – in batch-compatible fashion. Second, statistical validation strategies such as false-discovery rate (FDR) estimation are not implemented. This leads to unreliable sequence assignments and subjective manual assessment of spectra for validation. Third, existing solutions do not tie into any larger analytical framework, making integration with other (e.g. quantitative) data difficult. In contrast, shotgun proteomics has been sequencing peptides reliably for many years, and the inference, identification and quantification of proteins from constituent peptides has been automated to such a degree that the field has matured into answering biological questions at a more fundamental level^23^.

To fill this fundamental gap, we developed a fast, scalable database-matching tool called NucleicAcidSearchEngine (NASE) for the identification of RNA oligonucleotide tandem mass spectra. We implemented this new software within the OpenMS framework, an open-source toolset for processing mass spectrometric data^24^. NASE will be fully integrated into the primary distribution of OpenMS in the upcoming version 2.5, and will then be available for download as part of OpenMS at https://www.openms.de. In the meantime OpenMS builds containing NASE are available at https://www.openms.de/nase. Beyond speed and sensitivity, NASE provides advanced features like FDR estimation, precursor mass correction, and support for salt adducts. Powerful visualization capabilities are available through OpenMS’ data viewer. By supporting the common interface of The OpenMS Proteomics Pipeline^25^, NASE can be easily used in automated data analysis workflows. This interoperability also enables the label-free quantification of RNA oligonucleotides based on NASE search results – as a first in the field of nucleic acid mass spectrometry.

Based on three original datasets we demonstrate the capability of NASE to reliably identify a variety of RNA types from different sources, and show how data visualization and label-free quantification can augment the interpretation of identification results.

## Results

### RNA oligonucleotide MS datasets

Using nanoflow ion-pair liquid chromatography coupled to high-resolution tandem mass spectrometry (nLC-MS/MS), we generated three datasets from RNA samples of increasing complexity. First, oligonucleotides with the sequence of mature Drosophila let-7 microRNA, 21 nt in length, were produced synthetically in unmodified and modified (2’-O-methylated at the 3’ uridine) forms (“synthetic miRNA” dataset). We characterized a 1:1 mixture of both forms of this RNA. Replicate measurements were acquired using different normalized collision energy (NCE) settings in the mass spectrometer. Second, we prepared two samples of an *in vitro*-transcribed yeast lncRNA (NME1, 340 nt long), one of which was treated with an RNA methyltransferase (NCL1) catalyzing the 5-methylcytidine (m5C) modification (“NME1” dataset). These samples were subsequently digested with an RNA endonuclease (RNase) to generate oligonucleotide sequences of a length amenable to mass spectrometry. Third, we generated a sample of human total tRNA from a cellular extract – a complex mixture of highly modified RNAs (“human tRNA” dataset). This sample was again RNase-treated prior to nLC-MS/MS analysis.

### A powerful new search engine for RNA MS data

We developed a sequence database search engine for the identification of (modified) RNA sequences based on tandem mass spectra. The software, termed NucleicAcidSearchEngine (NASE), was implemented within the OpenMS framework and combines existing functionality (e.g. for data input/output, filtering, and FDR estimation) with newly developed features. (See Methods section for details.) Given a mass spectrometry data file and a FASTA file containing target and decoy (shuffled or reversed) RNA sequences as inputs, NASE generates oligonucleotide-spectrum matches with statistically meaningful FDR scores. OpenMS’ interactive viewer, TOPPView^26^, was extended to support RNA identification results obtained using NASE, mirroring and augmenting existing functionality for visualizing peptide identifications in proteomics experiments.

In addition to the built-in FDR calculation, NASE provides other features that set it apart from alternative tools that are currently available. Even with extensive preparation, nucleotide samples frequently contain salt adducts (in the form of cations attached to the phosphate backbone). NASE searches can take this into account, by allowing users to specify chemical formulas of adducts to consider in the precursor mass comparisons.

Furthermore, NASE supports the correction of precursor masses for MS2 spectra that were sampled from isotopologue peaks other than the monoisotopic one. Especially for longer sequences, MS2 precursor ions are often picked from higher-intensity, heavier isotopologues by the mass spectrometer’s data-dependent acquisition software. Without adjustment, the MS2 precursor masses would not closely match the theoretical (monoisotopic) masses of the correct oligonucleotides, leading to no assignment or incorrect matches. This feature thus greatly increases NASE’s ability to identify oligonucleotides with longer sequences.

Finally, through the OpenMS toolbox NASE enables seamless label-free quantification of the oligonucleotides that were identified in a sample. A corresponding analysis pipeline can be easily created and run using a graphical workflow editor. Supplementary Fig. 1 shows an example pipeline from our analysis of the NME1 data, using the editor that is conveniently included with OpenMS^27^.

### MS-based sequencing of an intact synthetic microRNA

In our analysis of data from the synthetic miRNA sample, we found a strong dependence of sequence coverage on the Normalized Collision Energy (NCE) value. Identical samples were run with NCE ranging from 5 to 55. The best results were obtained for an NCE of 20 (Supplementary Fig. 2). Subsequent LC-MS/MS analyses, including of the NME1 and tRNA samples, were thus carried out with this NCE setting.

At the optimal NCE, both unmodified and modified RNA were detected, and the location of the modification could be determined with high confidence. 874 spectra were identified that passed our hyperscore cutoff, matching sequences of length 5–21 nt, including the full-length let-7. The shorter sequences correspond to artefacts of incomplete solid-phase RNA synthesis, which are easily detectable by LC-MS. In the full 21-nt sequence we averaged over two-fold MS2 ion coverage of the let-7 sequence, with one or more forward (a-B/a/b/c/d) ion and one or more reverse (w/x/y/z) ion detected at each base (see Fig. 1, ion naming scheme from McLuckey et al.^28^). This demonstrates our ability to sequence even relatively long (>20 nt) RNAs.

**Figure 1:**
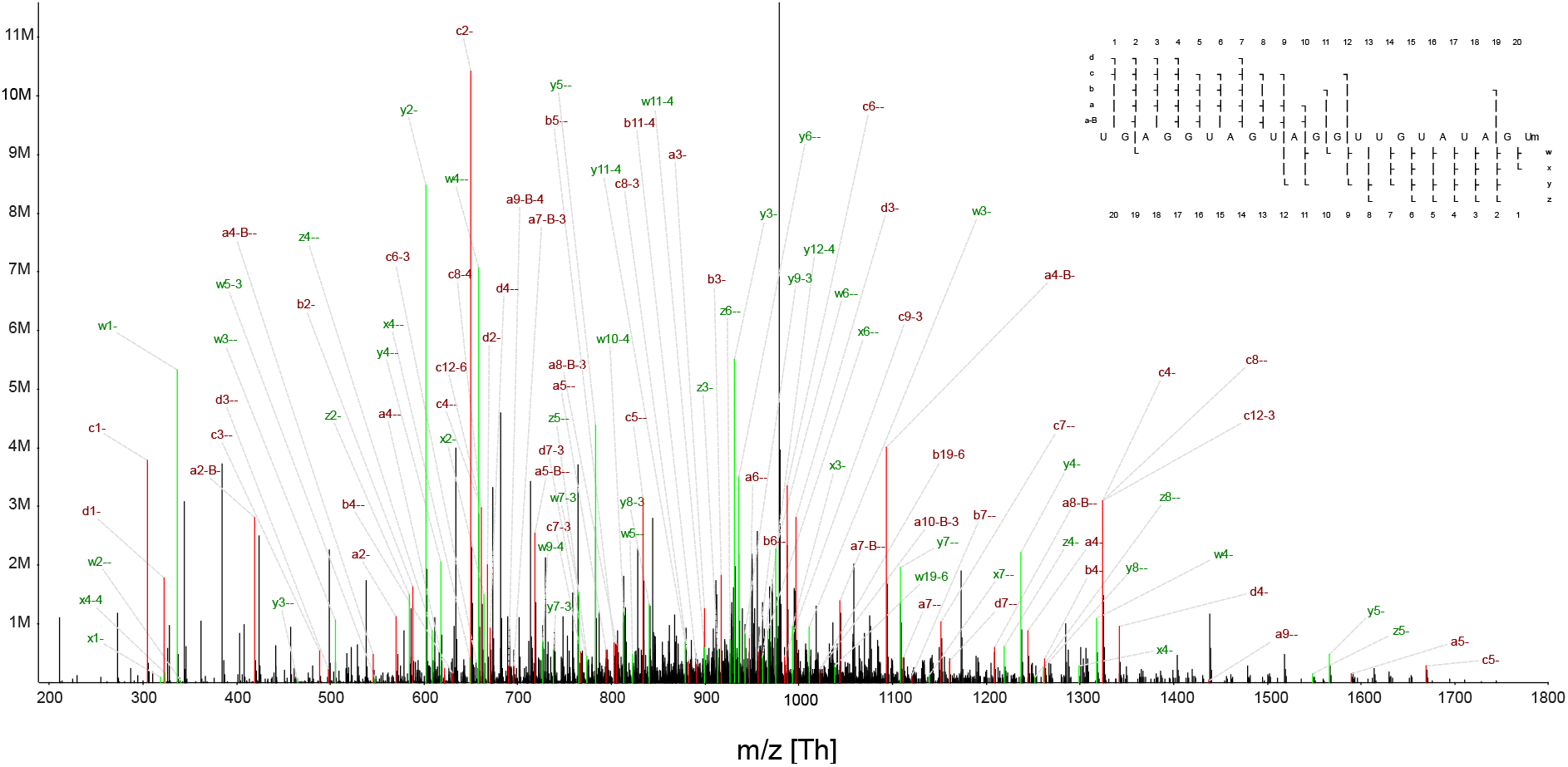
A tandem mass spectrum of synthetic let-7 denoting all of the assigned peaks. The primary ion was deprotonated seven times to give a charge state of –7 (m/z 971.55). The ion coverage plot in the upper right shows coverage for nine different types of fragment ion.

### Performance comparison of search engines for RNA MS data

We processed the NME1 data using the three search engines Ariadne, RNAModMapper, and NASE. We ran target/decoy database searches using m5C as a variable modification and compared the results in terms of: A, the number of identified spectra at different FDR thresholds; B, the sequence length distribution of the identified oligonucleotides at 5% FDR (Fig. 2). NASE identified significantly more spectra at a given confidence level than the other tools. It also found longer oligonucleotides, which would be more informative for identifying RNAs in complex samples. About 8% of the oligonucleotide-spectrum matches generated by NASE at 1% FDR included sodium adducts and would have been missed without the adduct search capabilities.

**Figure 2:**
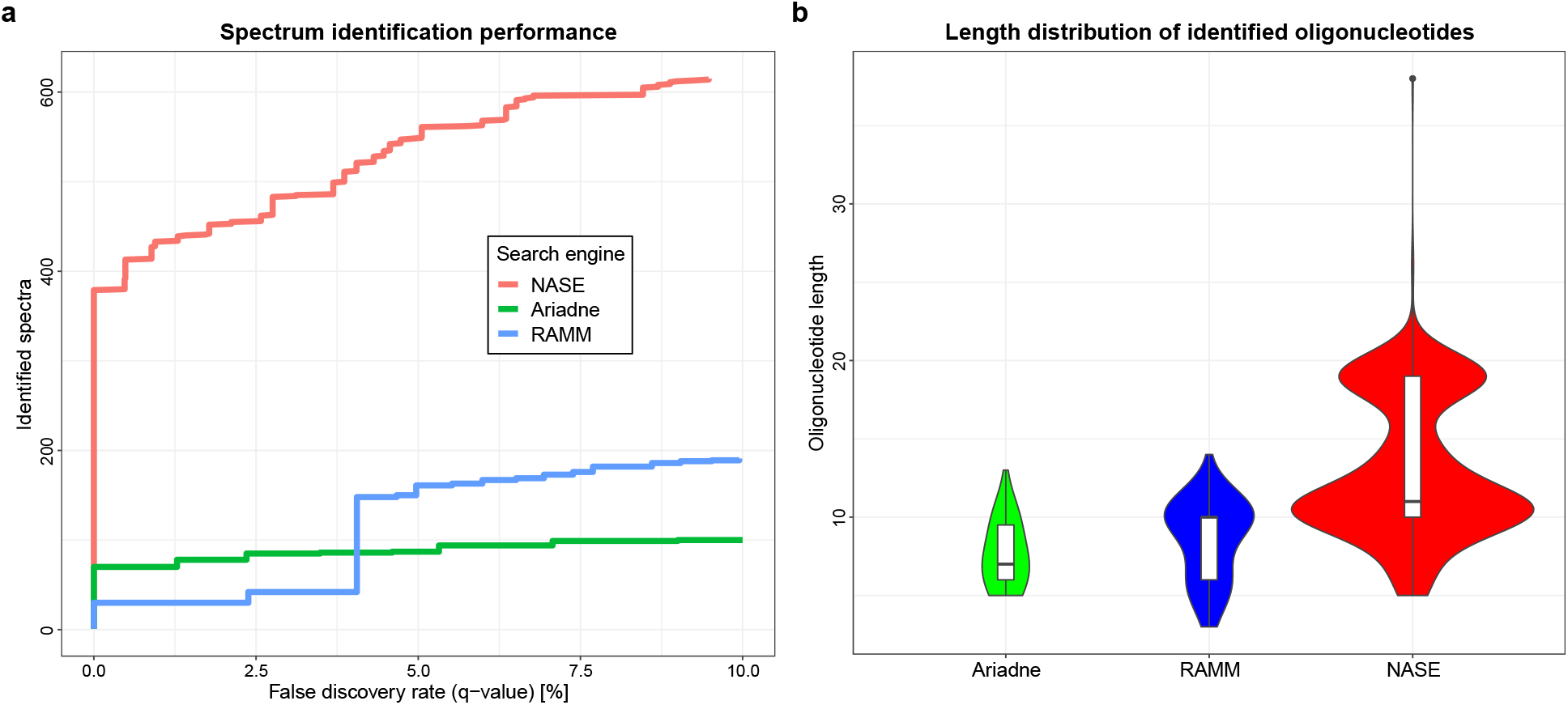
Performance comparison of RNA identification engines – Ariadne, RNAModMapper (RAMM) and NucleicAcidSearchEngine (NASE) – based on searches of the NME1 data. (a) Number of successfully identified spectra plotted against the q-value, a measure of the false discovery rate, which was calculated from a target/decoy database search using each of the three tools. (b) Sequence length distribution of identified oligonucleotides for each tool at a confidence level of 5% FDR.

Note that Ariadne’s performance in this comparison was hampered by the fact that a recommended tool for data preprocessing, the commercial software SpiceCmd, was not available to us. RNAModMapper had previously been evaluated based on searches against “expected” sequences only (i.e. no decoys), followed by manual validation of spectral assignments^22,29^.

### Detection of differential methylation sites in a benchmark sample

To assess the performance of our software at detecting RNA modifications, we compared the NASE search results for the NME1 lncRNA with and without NCL1 incubation (Fig. 3a). We considered results at a high confidence level of 1% FDR; at this level, 74% sequence coverage was achieved for both the control and the NCL1-treated sample. As Fig. 3a shows, there is good agreement between the unmodified oligonucleotides that were identified in both samples, indicating that our method works reproducibly. While a number of m5C-modified oligonucleotides were identified in the control sample, all except two of these false positives were observed in only a single oligonucleotide-spectrum match – in proteomic LC-MS/MS experiments, such “single hits” would be commonly filtered out^30^. We suspect that trace amounts of carry-over from earlier test runs of the NCL1 sample on the same chromatographic column may have caused these identifications in the control sample. Nonetheless, two modified oligonucleotides, “UCACAAAU[m5C]G” (at position 21–30 in the NME1 sequence) and “UAACC[m5C]AAUG” (position 299–308), were identified only in the NCL1-treated sample, based on 5 and 4 spectra in multiple charge states (−2 to –4 and –3 to –4, respectively). These (isobaric) oligonucleotides thus provide strong evidence for true m5C modification sites. Illustrating this, Fig. 4a shows a data section from the NCL1-treated sample, visualized as a two-dimensional LC-MS map. Identifications of the unmodified, adducted, and modified variants of the two oligonucleotides are displayed in the context of MS1 signal intensities. At the bottom, the isobaric oligonucleotides “UCACAAAUCGp” (left) and “UAACCCAAUGp” (right) can be seen eluting with slight separation. (In our notation, “p” at the end of a sequence represents the 3’ phosphate generated by RNase T1 cleavage.) In the middle, the corresponding mono-methylated oligonucleotides are convincingly detected, with a mass shift of 14 Da and a slight RT shift relative to their unmethylated counterparts. At the top, the oligonucleotide “UCACAAAUCGp” was identified with a sodium adduct (mass shift of 23 Da). A corresponding image showing the loss of signal for the modified oligonucleotides in the control sample is available as Supplementary Fig. 3. Fig. 4b compares spectrum matches for the two modified oligonucleotides, showcasing the high quality of the matches as well as our MS2 visualization capabilities, including the newly added ion coverage diagrams.

**Figure 3:**
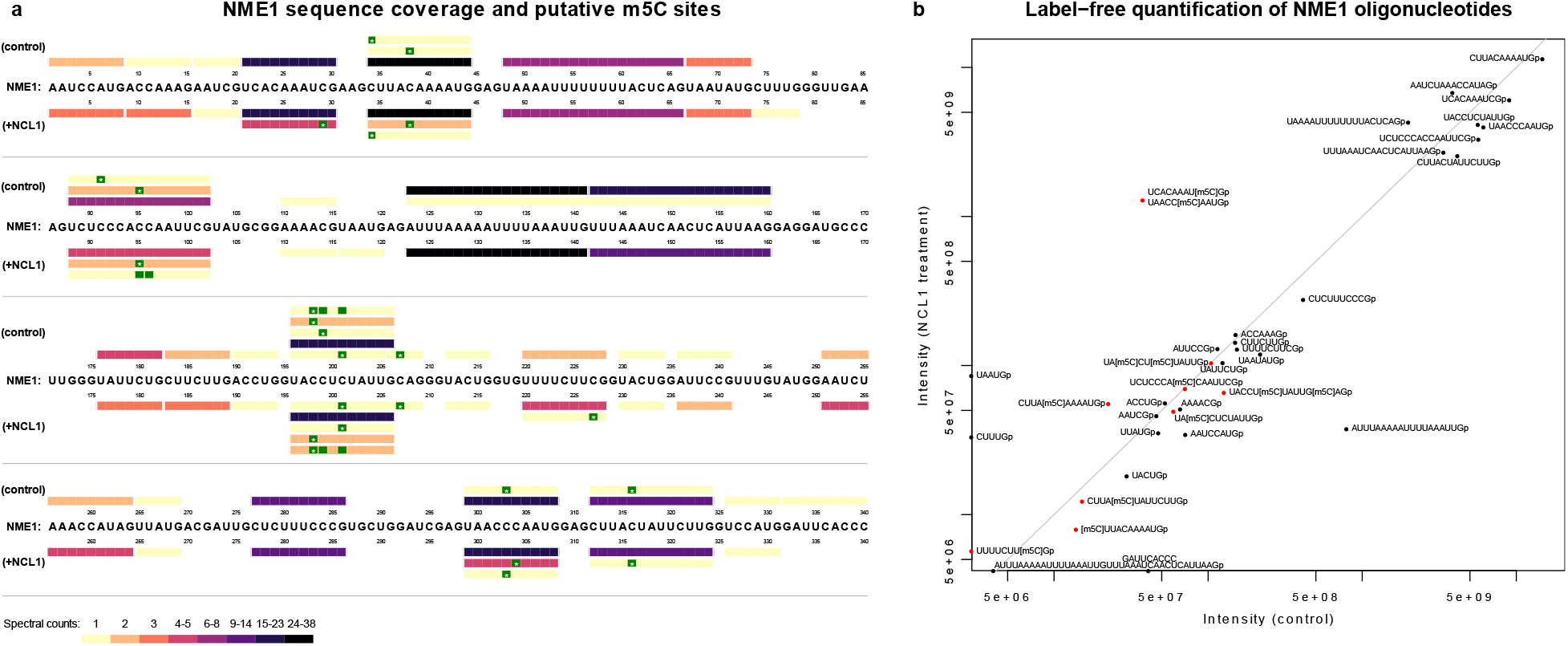
Comparison of the NME1 control and NCL1-treated sample based on NASE search results at 1% FDR. (a) Coverage plot showing oligonucleotides identified in the respective sample above/below the NME1 RNA sequence. Bars representing oligonucleotides are colored according to their number of identifications (spectral counts). Putative 5-methylcytidine (m5C) modification sites are marked in green. Sites with an asterisk (*) were uniquely localized, while “blank” sites indicate uncertainty between two possible locations, due to the absence of discriminating peaks in the corresponding mass spectrum. (b) Label-free quantification results for identified oligonucleotides, comparing feature-based signal intensities in the two samples. Intensities were aggregated over multiple charge and adduct states, where applicable. m5C-modified oligonucleotides are marked in red. Oligonucleotides that were quantified in only one of the samples are shown directly on the x and y axis, respectively. The grey diagonal line represents equal intensity in both samples.

**Figure 4:**
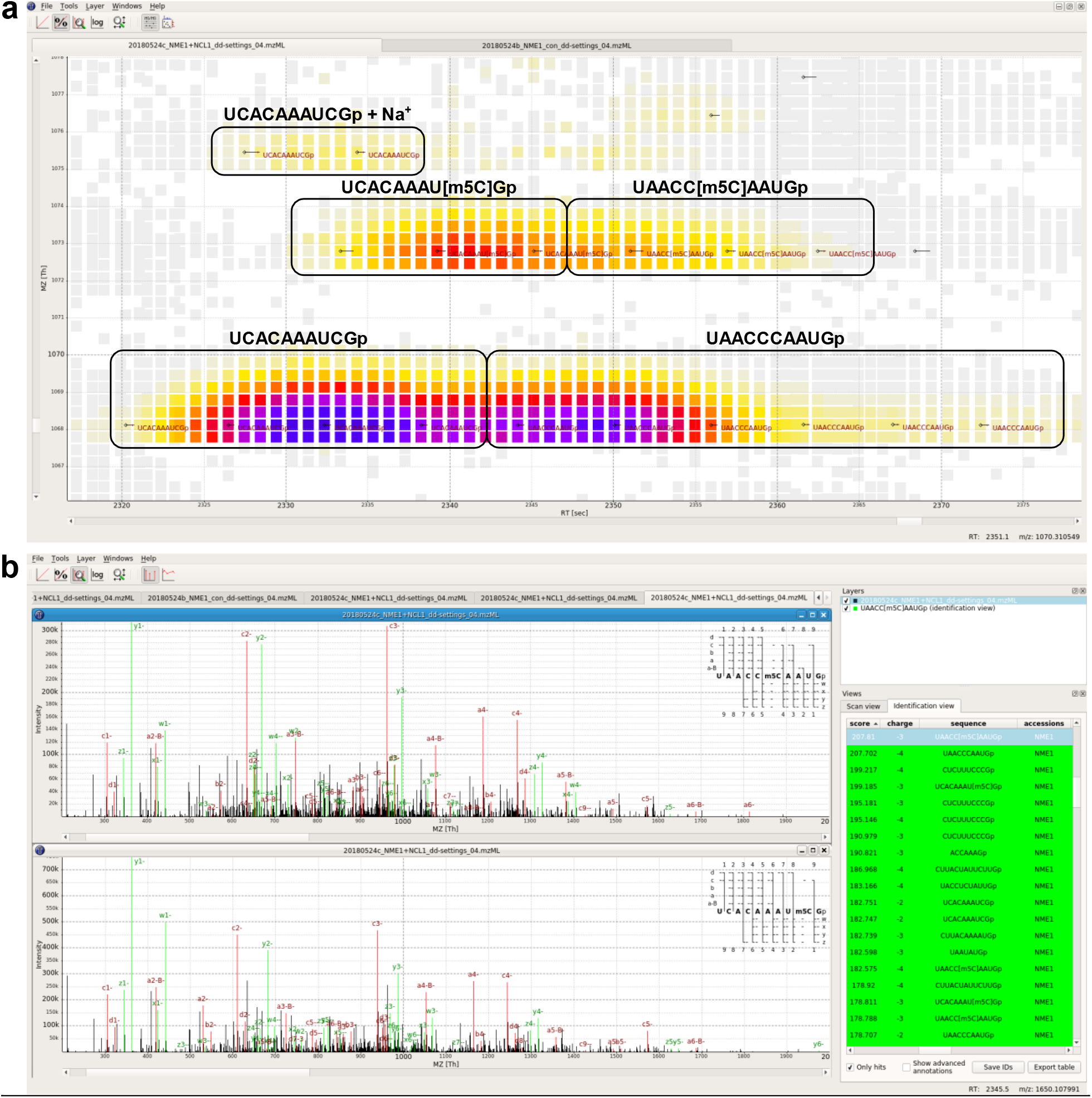
Screenshots from TOPPView showing data from the NCL1-treated NME1 sample. (a) MS1 view (RT-by-m/z) of a data section. LC-MS peaks are shown as small squares, colored according to their signal intensities. Small black diamonds and horizontal lines indicate MS2 fragmentation events; oligonucleotide sequences identified by NASE from the MS2 spectra are shown in dark red font. Rounded black boxes give an approximate outline of the chromatographic peaks corresponding to the oligonucleotides shown in black. These manual annotations are based on the identified sequences and signal intensity patterns. All oligonucleotides shown have a charge state of –3. (b) “Identification view” comparing two MS2 spectra, identified by NASE as the sequences “UAACC[m5C]AAUGp” and “UCACAAAU[m5C]Gp”. Matching peaks between the acquired and theoretical spectrum are annotated and highlighted in red and green. In the top-right corner of each spectrum plot, an ion coverage diagram shows which of the theoretical fragment ions of the sequence were matched in the MS2 spectrum (in any charge state).

### Label-free quantification of RNA MS data

We quantified the identified oligonucleotides in the two NME1 samples, using a label-free, feature detection-based approach. Fig. 3b summarizes the results. Although all oligonucleotides come from the same RNA, they were quantified with signal intensities spanning several orders of magnitude. This is indicative of widely varying ionization efficiencies during MS analysis, a common caveat that generally limits label-free quantification to relative comparisons between similar samples.

Of 37 and 40 oligonucleotides that were identified at 1% FDR in the control and NCL1-treated sample, respectively, 34 could be quantified in each sample (corresponding to 92% and 85% success rates). Most oligonucleotides were quantified at similar levels in both NME1 samples, with putative m5C-modified oligonucleotides generally found at lower intensities. The notable exception is the pair of modified oligonucleotides “UCACAAAU[m5C]G”/“UAACC[m5C]AAUG” already discussed above. While the chromatographic peaks for the unmodified oligonucleotides “UCACAAAUCG” and “UAACCCAAUG” were distinct enough to allow separate quantification of each, their modified variants could only be quantified together. The difference in signal intensities for these modified oligonucleotides between the control and NCL1-treated sample is clearly visible in Fig. 4a and Supplementary Fig. 3. This difference is exacerbated in the label-free analysis by the fact that only one corresponding identification was made in the control sample, while multiple charge states were identified, quantified and aggregated in the NCL1-treated sample. (The other obvious outlier, with the sequence “AUUUAAAAAUUUUAAAUUG”, was eluted at the very end of the chromatographic gradient and thus could not be quantified reliably.)

More advanced capabilities for LC-MS-based quantification, including retention time alignment, inference of identified analytes across samples, and labelling approaches, are already available in OpenMS for proteomics experiments. With future improvements to the support for nucleic acids in the framework, these features will become available for RNA analyses as well.

### Analysis of a complex, highly modified tRNA sample

Previous work on tRNA has shown that it is heavily modified^31^. Our analysis confirms this. We ran NASE on the “short RNA” fraction of a cell extract sample that had been digested with RNase T1. We searched for 26 variable modifications with different molecular masses, which had previously been identified to be present in yeast or human tRNA^32,33^. Most of these represent sets of isobaric modifications which we cannot distinguish, such as position-specific variants of the same modification; e.g. “m1A” was used to represent any singly-methylated adenosine (incl. Am, m6A etc.). Note that it was not feasible to search this dataset with this high number of variable modifications using other available database-matching tools (RNAModMapper, Ariadne). At an FDR cutoff of 5%, 1341 spectra were matched to 236 different oligonucleotide sequences. The sequences of human tRNAs are highly similar, especially for tRNAs of one isotype, i.e. tRNAs that bind the same amino acid. Consequently, only 38 (16%) of the identified oligonucleotides map to a unique tRNA sequence; however, 225 (95%) map uniquely to a single tRNA isotype. The sequence coverage, when counting all matching oligonucleotides for each of the detected tRNA sequences, ranged from 8.1% up to 54.8% (see Fig. 5a), with a median coverage of 20.8%. Coverage levels along the tRNA sequences were far from uniform, with the majority of identified oligonucleotides overlapping the anticodon loop and 3’ anticodon stem, or the T-loop and 3’ T-stem (Fig. 5b). We hypothesize that the corresponding parts of the tRNA structure are more amenable to RNase T1 digestion than other regions.

**Figure 5:**
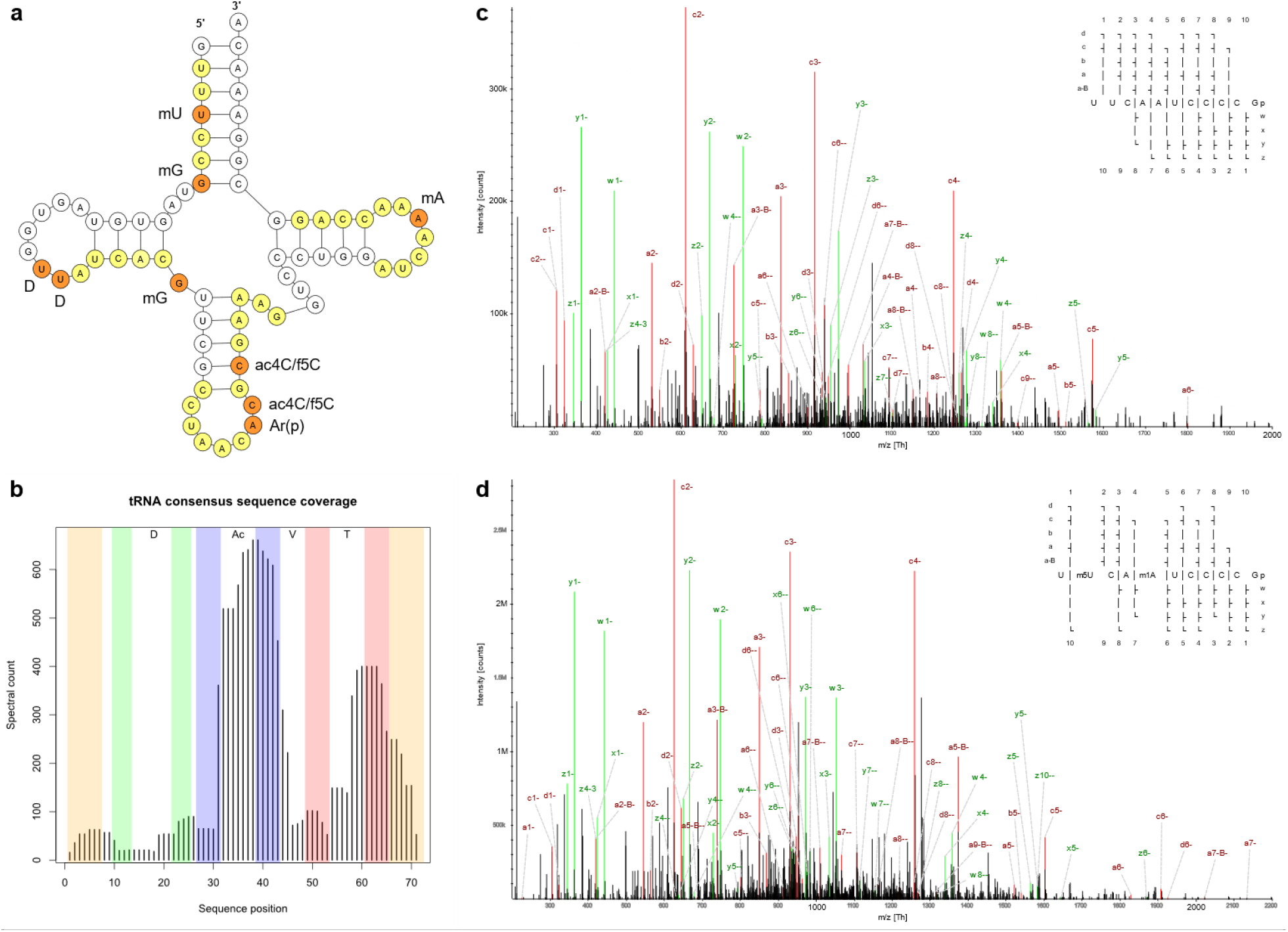
Human tRNA analysis results. (a) A schematic depiction of *Homo sapiens* tRNA-Val^AAC^-3–1. Sequences which we detected at 5% FDR are highlighted in yellow for unmodified, and orange for modified residues. Total coverage is 54.8%. The tRNAdb entry for tRNA-Val agrees with our findings, except for the methylation at U4 (based on four identified spectra) and the three modifications in the anticodon loop and stem (bottom right, based on two identified spectra). (b) Aggregated coverage of the consensus tRNA sequence by oligonucleotides identified in the human tRNA dataset. Every spectrum match at 5% FDR is counted. Some oligonucleotide positions in long tRNAs (tRNA-Leu, tRNA-SeC, tRNA-Ser) were adjusted to fit the consensus sequence. Complementary regions in the acceptor stem (orange), D-stem (green), anticodon stem (blue) and T-stem (red) are highlighted. D: D-loop, Ac: anticodon loop, V: variable region, T: T-loop. (c),(d) Tandem mass spectra of an oligonucleotide from human tRNA-Ala^AGC^, which was characterized with and without post-transcriptional modifications. Both primary ions shown here were triply-charged (m/z 1154.14 and 1163.48, respectively). In our analysis at 5% FDR, the modified and unmodified forms were both identified 13 times each, with charge states ranging from –2 to –5 and from –3 to –5, respectively.

Many of the oligonucleotides we identified contained multiple modifications. In the search, up to three modifications per oligonucleotide were allowed, to limit the combinatorial space of modified sequences that needed to be explored. Of the unique oligonucleotides identified at 5% FDR, 11% were unmodified (accounting for 16% of the identified spectra), while 36% carried one, 26% carried two, and 25% carried three modifications (accounting for 45%, 23% and 16% of the identified spectra, respectively).

All modifications included in the search, except queuosine, wybutosine and their derivatives, were detected as part of identified oligonucleotides. However, the prevalences of different modifications differed widely – see Table 1 for details.

**Table 1:** Summary of modifications detected in the HAP1 tRNA data using NASE at a 5% FDR level. Columns: 1. Short code of the modification specified as a search parameter. 2. The set of modifications implied by the corresponding mass shift, since e.g. position-specific variants of a modification (Am, m1A, m6A etc.) generally cannot be distinguished. 3. Number of identified oligonucleotide-spectrum matches with at least one instance of the corresponding modification in the sequence. 4. Number of unique oligonucleotides with at least one corresponding modification among the search results.

| Search mod.<br>(short code) | Represented isobaric modification(s) | Spectrum matches | Unique oligonucleotides |
| --- | --- | --- | --- |
| m1A | Adenosine monomethylation | 391 | 55 |
| m5U | Uridine or pseudouridine monomethylation | 247 | 46 |
| t6A | N6-threonylcarbamoyladenosine | 222 | 29 |
| m5C | Cytidine monomethylation | 173 | 24 |
| m2G | Guanosine monomethylation | 114 | 26 |
| D | Dihydrouridine | 107 | 26 |
| I | Inosine | 78 | 26 |
| acp3U | 3-(3-amino-3-carboxypropyl)uridine or -pseudouridine | 63 | 18 |
| i6A | N6-isopentenyladenosine | 55 | 13 |
| m2,2G | Guanosine dimethylation | 48 | 13 |
| ncm5s2U | 5-carbamoylmethyl-2-thiouridine | 43 | 17 |
| ac4C | N4-acetylcytidine or 5-formyl-2'-O-methylcytidine (f5Cm) | 37 | 14 |
| Ar(p) | 2'-O-ribosyladenosine (phosphate) | 26 | 14 |
| mnm5U | 5-methylaminomethyluridine | 23 | 5 |
| m1I | Inosine monomethylation | 20 | 7 |
| m1Im | Inosine dimethylation | 10 | 6 |
| f5C | 5-formylcytidine | 9 | 7 |
| io6A | N6-(cis-hydroxyisopentenyl)adenosine | 5 | 4 |
| m5Cm | Cytidine dimethylation | 3 | 3 |
| m5D | Dihydrouridine monomethylation | 2 | 2 |
| m6Am | Adenosine dimethylation | 2 | 1 |
| m5Um | Uridine or pseudouridine dimethylation | 1 | 1 |
| yW | Wybutosine | 0 | 0 |
| Q | Queuosine | 0 | 0 |
| o2yW | Peroxywybutosine | 0 | 0 |
| galQ | Galactosyl- or mannosyl-queuosine (manQ) | 0 | 0 |

Existing data on the modification landscape of human cytosolic tRNAs is incomplete (e.g. tRNAdb^33^ lists information for 20 genes covering 15 isotypes) and at least some modifications are differentially regulated, complicating comparisons. We will focus on cytosine monomethylation (mC, represented by m5C in our search) as one example that has been studied more thoroughly, e.g. via bisulfite sequencing to detect m5C. At 5% FDR we identified 24 unique oligonucleotides with unambiguous assignments of mC. The oligonucleotides contained one or two mC sites each and were supported by a total of 173 identified spectra. Each mC-containing oligonucleotide was associated with one unique or predominant (more matching genes) tRNA isotype. At the level of these isotypes, a total of 18 unique mC sites were identified. Seven of these sites agree with the “canonical” m5C sites in the VL junction of tRNAs at consensus sequence positions 48–50^7^. Cytosine methylation at position 34 in the anticodon, previously reported^34^ as m5C for tRNA-Leu^CAA^ and 2’-O-methylcytidine (Cm) for tRNA-Met in tRNAdb, was here observed for tRNA-Met and tRNA-Trp. Methylation at C32 was detected in several tRNAs (tRNA-Gln^CTG^, tRNA-Leu^TAA^, tRNA-Phe^GAA^, tRNA-Trp, tRNA-Val^CAC^); correspondingly, Cm is reported at this position for tRNA-Gln and tRNA-Phe in tRNAdb.

We find that our results generally recapitulate annotated modifications in tRNAdb, in regions where we have sequence coverage and with the caveat that we cannot distinguish between isobaric modifications (including uridine/pseudouridine). Known recurring modifications that we identify in several tRNAs include monomethylation at G10, mono-or dimethylation at G26, and monomethylation at A58. In many cases we find additional, alternatively modified (or unmodified) variants of “expected” oligonucleotides. In particular, for an oligonucleotide that matches the T-loop region in several tRNA-Ala genes we robustly detect the unmodified form and a doubly methylated form (mU55 and mA58; see Fig. 5c/d for annotated spectra). The ability to identify and localize multiple different modifications simultaneously is a unique advantage of the oligonucleotide MS approach. In this example, each form was matched to 13 different spectra at our 5% FDR cutoff; singly methylated variants were also detected in lower numbers. For the equivalent oligonucleotides in tRNA-Cys and tRNA-Gln (or tRNA-Thr^TGT^ – same partial sequence), we found multiple matches of both the double methylation and a single methylation at A58. In oligonucleotides overlapping the anticodon loop and the 3’ anticodon stem, we detect multiple forms for tRNA-Gly^CCC^ (mU39 with and without mU32), tRNA-Met (either or both of mC34 and t6A37), tRNA-Val^CAC^ (unmodified and mC32) and tRNA-iMet (unmodified and t6A38). For tRNA-Ser we observe several different forms at this location – primarily mA37 and mU44 with or without N6-threonylcarbamoyladenosine (t6A) at A42 for tRNA-Ser^GCT^, and N6-isopentenyladenosine (i6A) at A37 with either or both of mU39 and mU44 for tRNA-Ser^*GA^. Based on our data it is impossible to determine whether these and other cases correspond to partial modifications of a particular tRNA, or to mixtures of differently modified tRNAs from separate genes. However, overall these results support newer findings that question the stoichiometric and static nature of tRNA modifications, and favor the notion of a complex and dynamic tRNA modification landscape^35^.

## Discussion

NASE is a new open-source database search engine for RNA, optimized for high-resolution MS data. It supports arbitrary modifications, salt adducts, and FDR estimation through a target/decoy search strategy. Moreover, integration with the OpenMS toolbox enables high-quality data visualization, e.g. for manual validation of spectral assignments, and label-free quantification of RNA oligonucleotides. We have tested NASE against a range of sample types and complexities, spanning synthetic nucleic acids, *in vitro*-transcribed sequences, and cell extracts. In all of these experiments we have been able to effectively identify RNA sequences and their modifications.

NASE contains many unique functionalities that are not currently realized in other database search tools for RNA. To our knowledge, no other tools account for precursor mass defect resulting from instrumental selection of higher isotopologue peaks. This functionality is a major contributor to the excellent performance of NASE in identifying longer oligonucleotides compared to other database-matching tools. NASE also provides powerful correction for cation adduction events, which lessens the impact of sodium and potassium ions on sequence characterisation. In addition, OpenMS in general and NASE specifically were designed to be fast. Our search times for complex samples are orders of magnitude faster than other tools. The searches on the NME1 and let-7 data take seconds; the much more complicated 26-modifications search of the tRNA dataset took 24 hours in multithreaded mode (20 parallel threads) on our server. For comparison, an analogous search using RNAModMapper was not feasible, with an estimated running time of one month. An equivalent search with Ariadne did not return any modified oligonucleotides.

The open-source nature of OpenMS and NASE enables users to modify the software to fit their specific needs, to extend the existing functionality, and to create new interoperating programs. Already, many analysis tools have been implemented within the OpenMS framework to support mass spectrometry-based proteomics and metabolomics experiments. The present work, and here in particular the pioneering application of label-free quantification, gives a foretaste of the power of leveraging these methods for the analysis of nucleic acid data. Future developments will streamline the use of OpenMS tools and algorithms, e.g. for improved quantification and comparisons across many samples, in the field of epitranscriptomics. In conclusion, the development of NASE is an important step towards the large-scale analysis of RNA by mass spectrometry.

## Methods

### Liquid chromatography-tandem mass spectrometry

RNA samples were separated by reversed-phase ion-pair liquid chromatography (using 200 mM HFIP + 8.5 mM TEA in H_2_O as eluent A, and 100 mM HFIP + 4.25 mM TEA in methanol as eluent B) and characterized by negative ion MS/MS in a hybrid quadrupole-orbitrap mass spectrometer (Q Exactive HF, Thermo Fisher). A gradient of 2.5 to 25% eluent B eluted oligonucleotides from various lengths of nanoflow Acclaim PepMap C18 solid phase (Thermo Fisher) at 200 nL/min. The length of gradient was varied according to the complexity of the sample. Precursor ion spectra were collected at a scan range of 600 to 3500 m/z at 120k resolution in data-dependent mode, with the top five MS1 species selected for fragmentation and MS2 at 60k resolution.

### RNA samples

A variety of RNA samples were characterized by nanoflow LC-MS/MS (nLC-MS/MS) and sequence analysis performed using NASE. Initial work was carried out on a mature *Drosophila* let-7 sequence that was prepared by solid-phase synthesis and purchased from IDT. This sequence is a 21 nt long microRNA that was among the first miRNAs to be characterized^36^. The RNA was chemically synthesized in unmethylated and methylated forms, i.e. with or without a 2’-O-methyluridine (Um) at position 21. A sample was prepared by mixing both forms, and was characterized by nLC-MS/MS without further processing, but with varying normalized collision energy (NCE) settings to give different levels of precursor fragmentation.

Subsequent experiments were carried out on NME1, a 340 nt long *Saccharomyces* lncRNA. NME1 RNA was generated by *in vitro* transcription, and two samples with and without NCL1 enzyme treatment were prepared. NCL1 is a yeast RNA methyltransferase that catalyzes the 5-methylcytidine (m5C) modification^37^. RNA was extracted and digested with RNase T1 prior to nLC-MS/MS. This endonuclease generates shorter oligonucleotides by cleaving immediately after guanosine residues.

The most complex sample was a solution of digested crude human cellular tRNA, which was isolated from HAP1 tissue culture using an RNeasy kit (Qiagen) as according to the manufacturer’s instructions. Briefly, RNAs can be fractionated by length by differential elution, with RNAs less than 200 nucleotides mostly made up of tRNA, and the larger fraction being mostly rRNA. The “shorter” RNA fraction was digested with RNase T1, and the resultant oligonucleotides were characterized by nLC-MS/MS.

### NucleicAcidSearchEngine implementation

NASE was implemented in C++ within the OpenMS framework. The OpenMS library was extended with classes representing (modified) ribonucleotides (based on data from the MODOMICS database^38^), RNA sequences, and riboendonucleases. A new generalized data structure for spectrum identification results (supporting peptides/proteins, nucleic acid sequences, and small molecules) and an algorithm for theoretical spectrum generation of RNAs were added as well. NASE itself is a new executable tool that supports the common interface of The OpenMS Proteomics Pipeline^25^.

Data processing with NASE works as follows: Inputs are an RNA sequence database (FASTA format) and a mass spectrometry data file (mzML format). RNA sequences are digested in silico using enzyme-specific cleavage rules for the user-specified RNase. Tandem mass spectra are pre-processed (intensity filtering, deisotoping) and mapped to oligonucleotides based on precursor masses. Mass offsets due to salt adducts or precursor selection from heavier isotopologue peaks can be taken into account. Next, theoretical spectra of relevant oligonucleotides in the appropriate charge states are generated and compared to the experimental spectra; matches are scored using a variant of the hyperscore algorithm^39^. If the sequence database contains decoy entries, the resulting oligonucleotide-spectrum matches can be statistically validated through the automatic calculation of q-values, a measure of the FDR^40^. Supported output formats are an mzTab-like^41^ text file, suitable for further analysis, and an XML file, suitable for visualization in TOPPView.

In order to provide support for label-free quantification of identified oligonucleotides, NASE interfaces with the OpenMS tool FeatureFinderMetaboIdent (FFMetId). FFMetId handles the core step of the quantitative workflow, the detection of chromatographic features in the LC-MS data. As a variant of the proteomics tool FeatureFinderIdentification^42^, FFMetId provides targeted feature detection for arbitrary chemical compounds. NASE can write an output file with all relevant information about the oligonucleotides it identified, which is directly suitable as an input file for FFMetId.

### Data processing

#### Sequence database searches

For NASE analyses, all proprietary raw files were converted to mzML format^43^ without compression and with vendor peak-picking using MSConvert^44^ (https://github.com/ProteoWizard). The full list of fragment ion types (a-B, a, b, c, d, w, x, y, z) was considered for peak matching. Precursor and fragment mass tolerance were both set to 3 ppm. For precursor mass correction, the monoisotopic up to the fifth (+4 neutrons) isotopologue peak were considered.

The synthetic let-7 data was searched with NASE using unspecific cleavage to account for incomplete RNA synthesis products. An extensive set of potential adducts (Na^+^, K^+^, Na_2_^2+^, K_2_^2+^, NaK^2+^, Na_3_^3+^, K_3_^3+^, Na_2_K^3+^, NaK_2_^3+^) was used because of the substantial salt that remained from the RNA synthesis reactions. Two copies of the let-7 sequence, one with a fixed 2’O-methylation of uridine (Um) at the 5’ position, were specified in the FASTA sequence file. The small size of the sequence database prevented the use of a target/decoy approach for FDR estimation. We thus used a stringent hyperscore cutoff of 150 (corresponding to the 1% FDR in the tRNA sample, see below) to define a high-confidence set of results.

The NME1 data analysis used RNase T1 digestion with one allowed missed cleavage. m5C was set as a variable modification; up to two modifications per oligonucleotide were considered. Na^+^ was specified as a potential adduct. The sequence database contained the NME1 (target) sequence as well as a shuffled decoy sequence.

In our search of the tRNA data, 26 variable modifications (based on previous findings in yeast and human tRNA) were specified, at a maximum of three modifications per oligonucleotide. See Table 1 for the full list of modifications. Na^+^ was specified as a potential adduct. The FASTA file contained 420 human tRNA sequences collected from the tRNA sequence database tRNAdb^33^ (http://trna.bioinf.uni-leipzig.de) plus the same number of reversed decoy sequences. The digestion parameters were set to RNase T1 with up to two missed cleavages.

#### Search engine comparison

The NME1 data was processed with two other publicly available RNA identification engines, in addition to NASE: Ariadne^20^ and RNAModMapper^22^. To this end, the raw files were converted to MGF format using MSConvert. Cleavage and variable modification settings in the searches were the same as for NASE and appropriate for the samples.

For Ariadne, the online version at http://ariadne.riken.jp was used in October 2018. The “Calc as partial modifications” option was enabled. The precursor and fragment mass tolerances were left at their default values (5 and 20 ppm). Alternatively, using the parameters from the Taoka et al.^45^ (20 and 50 ppm) made no appreciable difference for Ariadne’s performance in our tests.

For RNAModMapper, a program version from July 2018 was used with settings recommended by the author, Ningxi Yu. All available ion types (a-B, w, c, y) were enabled; precursor and fragment mass tolerance were set to 0.02 and 0.1 Da, respectively.

#### Label-free quantification

In order to perform label-free quantification on the NME1 dataset, target coordinates (chemical sum formulas, charge states, median retention times) for oligonucleotides identified at 1% FDR were exported from NASE.

Based on these coordinates, feature detection in the LC-MS raw data (mzML files) was carried out with the OpenMS tool FeatureFinderMetaboIdent. The results were exported to text format using OpenMS’ TextExporter, for subsequent processing and visualization in R 3.5.1^46^. Results from both NME1 samples were merged and feature intensities for oligonucleotides were summed up over multiple charge and adduct states, where available. To ensure comparability, manual adjustments were made in a few cases where modified oligonucleotides had been identified with different m5C localizations in the two samples.

## Acknowledgements

We would like to thank the following people: Lina Vasiliauskaitė for preparing the HAP1 tRNA sample. Ningxi Yu for suggesting optimal parameters for running his RNAModMapper tool. All contributors to OpenMS; especially Hannes Röst for his efforts and useful feedback during the code review for this project. BAG acknowledges funding from NIH grant GM110174 and a UPenn Epigenetics Institute Pilot grant.

## Author Contributions

SW, TS and HW developed the software. SW, BA and HW analyzed the data and wrote the manuscript. BA performed the LC-MS/MS experiments. HSR prepared the NME1 samples. OK, TK and BAG provided resources and high-level supervision. All authors read and approved the manuscript.

## Data Availability

Mass spectrometry data files and search results (as well as label-free quantification results for the “NME1” dataset) were deposited in the PRIDE^47^ repository with dataset identifiers PXD012094 (synthetic let-7), PXD012095 (NME1) and PXD012097 (human tRNA).

## Supplementary Figures

**Supplementary Figure 1:**
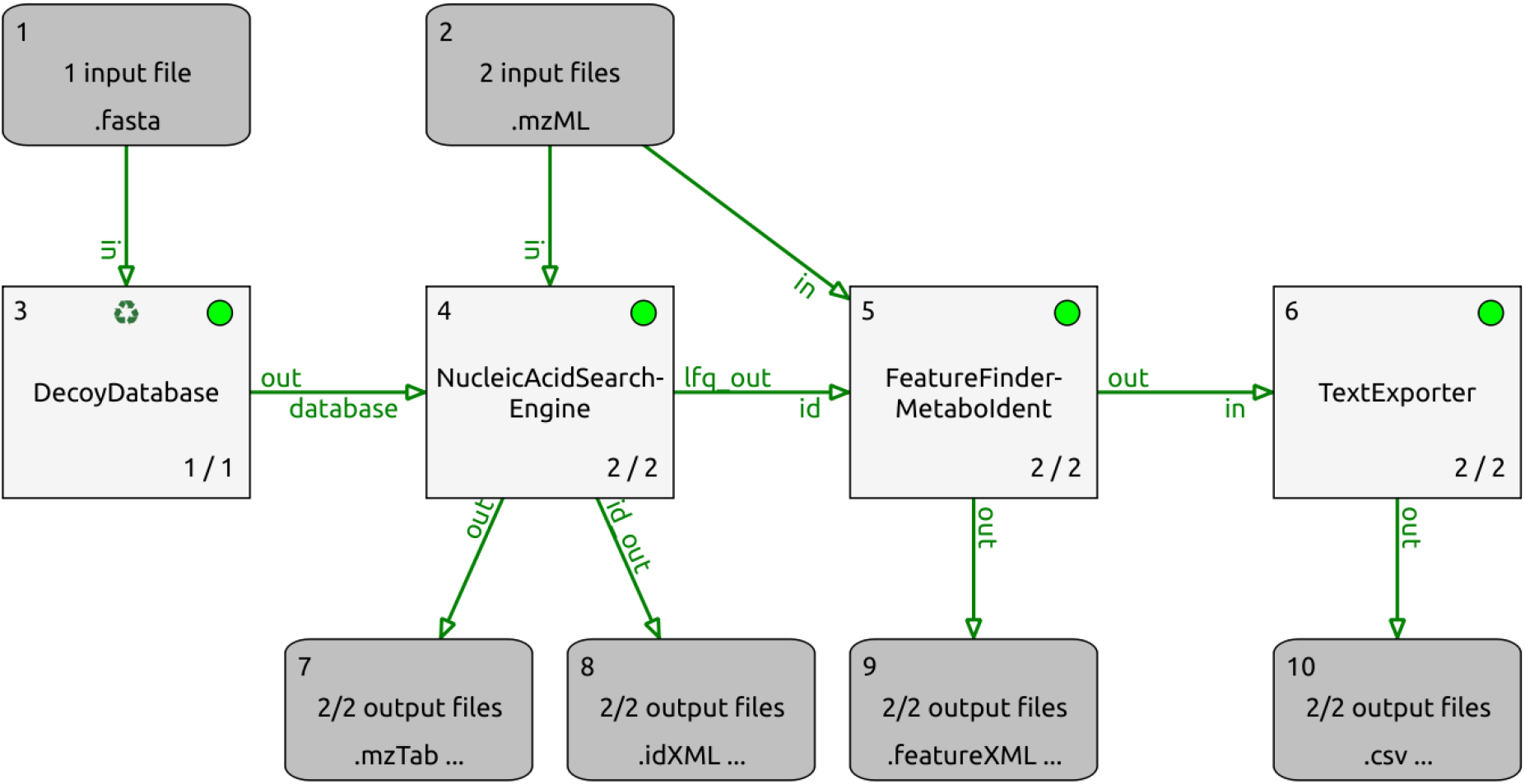
Data analysis pipeline for the NME1 data, comprising target/decoy database generation, database search (incl. FDR estimation and filtering), targeted feature detection and data export. Screenshot from TOPPAS, the OpenMS workflow editor. The whole pipeline ran in only 12 seconds (single-threaded) on our server.

**Supplementary Figure 2:**
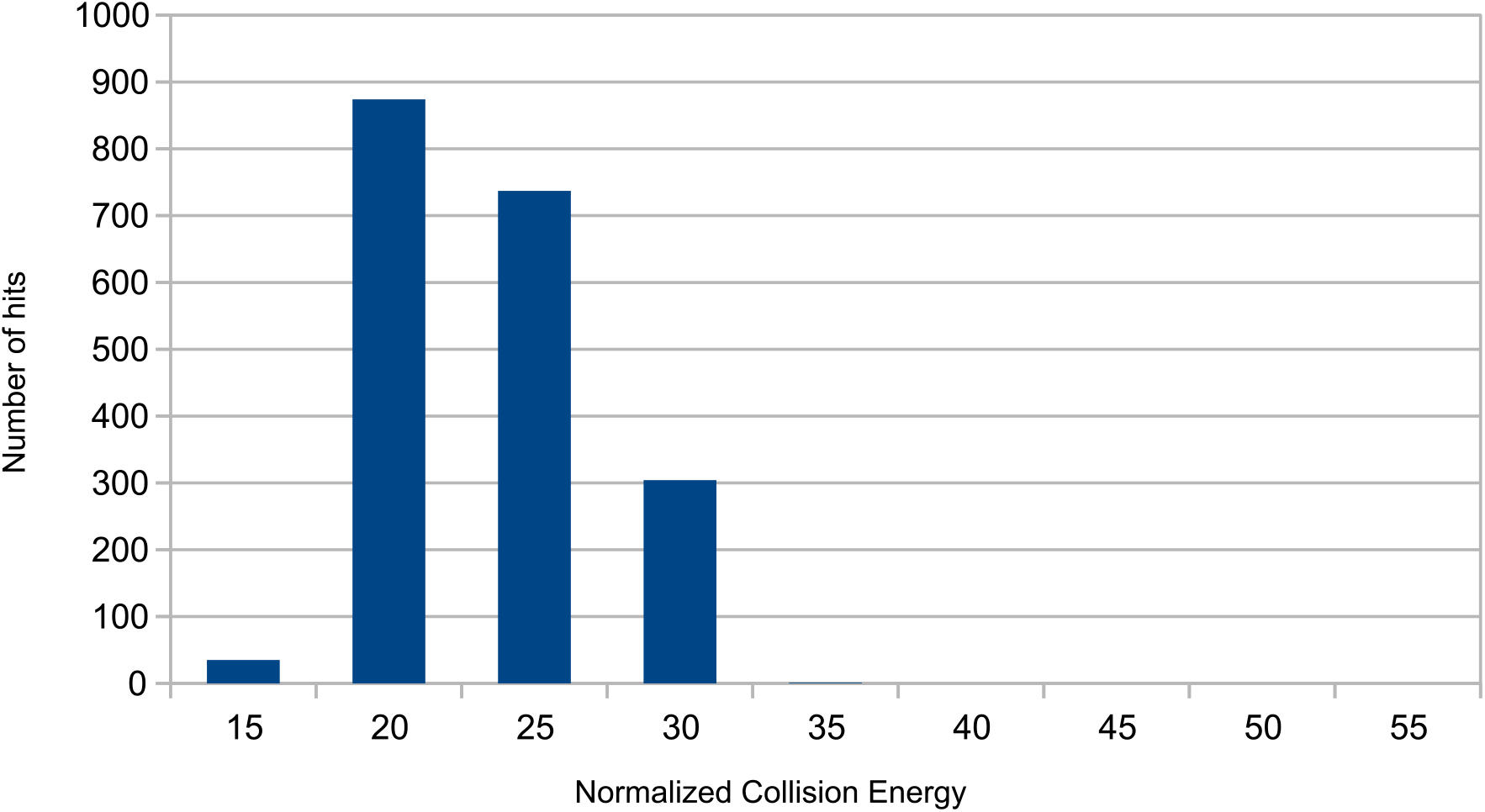
Graph showing the relationship between Normalized Collision Energy (NCE) used in HCD and the number of NASE search hits scoring above our cutoff in replicate runs of the let-7 sample. The optimum NCE is 20.

**Supplementary Figure 3:**
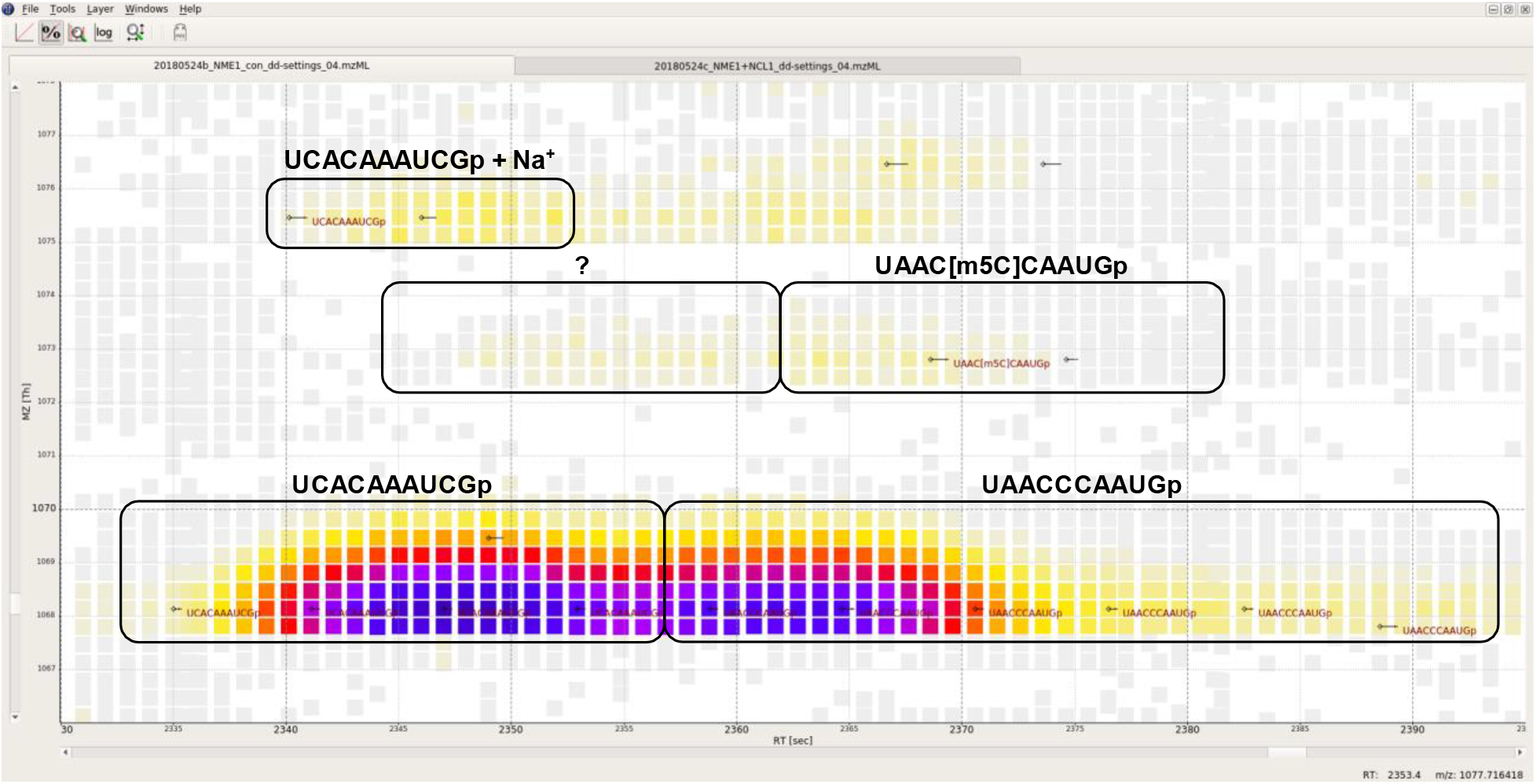
Annotated screenshot from TOPPView showing data from the NME1 control sample, corresponding to the NCL1-treated data shown in Fig. 4a. Note the loss of signal intensity and sequence identifications for the methylated oligonucleotides, compared to Fig. 4a. Due to a lower-quality MS2 spectrum, the m5C site in “UAACCCAUGp” has here been localized to the second, not third cytidine.

